# Sex ratio and sexual conflict in a collective action problem

**DOI:** 10.1101/2020.01.29.924316

**Authors:** C. Lindstedt, N. Gerber, H. Kokko

**Author notes:** Corresponding author: Carita Lindstedt, Department of Biological and Environmental Sciences, University of Jyväskylä, P.O. Box 35, FI-40014 University of Jyväskylä, Finland.

## Abstract

The maintenance of cooperation is difficult whenever collective action problems are vulnerable to freeriding (reaping the benefits without contributing to the maintenance of the good). We identify a novel factor that can make a system tolerate an extent of freeriding. If a population consists of discrete types with demographically distinct roles, such that the success of one type does not imply it can spread to replace other types in the population, then collective goods may persist in the presence of free-riders because they are necessarily kept in a minority role. Biased sex ratios (e.g. in haplodiploids) create conditions where individuals of one sex are a minority. We show that this can make the less common sex contribute less to a public good in a setting where the relevant life-history stage — larval group defence against predators — does not feature any current breeding opportunities that might lead to confounding reasons behind sex-specific behaviour. We test our model with haplodiploid pine sawfly larvae, showing that female larvae are the main contributors to building the antipredator defence against predators.

**Significance statement:** Individuals in groups can cooperate to achieve something together, but with an evolutionary difficulty: if benefits of cooperation are shared equally among all, freeriders get the same benefit as others while paying less for it. We propose a novel reason why freeriding does not automatically spread until the collectively beneficial outcome is destroyed: sometimes groups consist of individuals of distinct categories, limiting freerider spread. If, for example, there are always fewer males than females, then even if every male becomes a freerider, the whole group still survives simply because not everyone can be male. Pine sawfly larvae defend against predators by regurgitating sticky fluids, but females contribute more to this common defence, and we show this example fits our model.

## Introduction

Group living organisms often face collective action problems, where cooperative individuals contribute to a good that can be exploited by many or all individuals living in the same group. This brings about problems of freeriding, and consequently of maintaining the collective good as a whole. If contributing is costly, and benefits are shared, it is not *a priori* guaranteed that contributing leads to higher fitness than reaping the benefit without having contributed to its production (1–5). One relevant factor is heterogeneity such that individuals vary in their costs and benefits of producing and taking advantage of the good (reviewed by (6)). For example, in many primates with strong dominance hierarchy, high-ranking individuals often contribute more to a collective good, despite not necessarily gaining as much net benefit from this behaviour (6). This differs from the expectations for homogenous group structures, where costs of contributing to the collective action and benefits gained from it are assumed to be equal for all group members.

Here we focus on a neglected possibility within the concept of heterogeneity: individuals may belong to demographically distinct categories with an inheritance system that prevents freeriding from automatically spreading across category boundaries. Freeriding may then evolve to be prevalent in the minority category without threatening the collective good as a whole. We first illustrate this principle with a toy model, then with a case of haplodiploid sex determination, where the categories are equivalent to males and females, and sex ratios are often biased. We use the full model to explain findings of sex-specific contribution to a collective good (group defence against predators) in pine sawfly larvae.

Our toy model (see Supplementary material for mathematical detail) considers the following scenario: Groups of five members are formed by sampling members from two distinct (but geographically overlapping) populations, such that population M (the majority) always contributes 3 members, while population m (the minority) always contributes 2 members. M-individuals contribute towards the common good with probability *x* or freeride (fail to contribute) with probability 1–*x*; for m-individuals the corresponding probabilities are *y* and 1–*y*. The group survives if more than half of its members contributed; among survivors, those that did not contribute enjoy higher fitness (contributing was costly) by a factor α. Intuition suggests that m-individuals might be more prone to freeride because group survival is possible even if freeriding becomes fixed in this population, as m-individuals never form large enough proportion of groups to alone cause the collapse of the good. This intuition is confirmed by mathematical analysis (Fig. 1).

**Figure 1.**
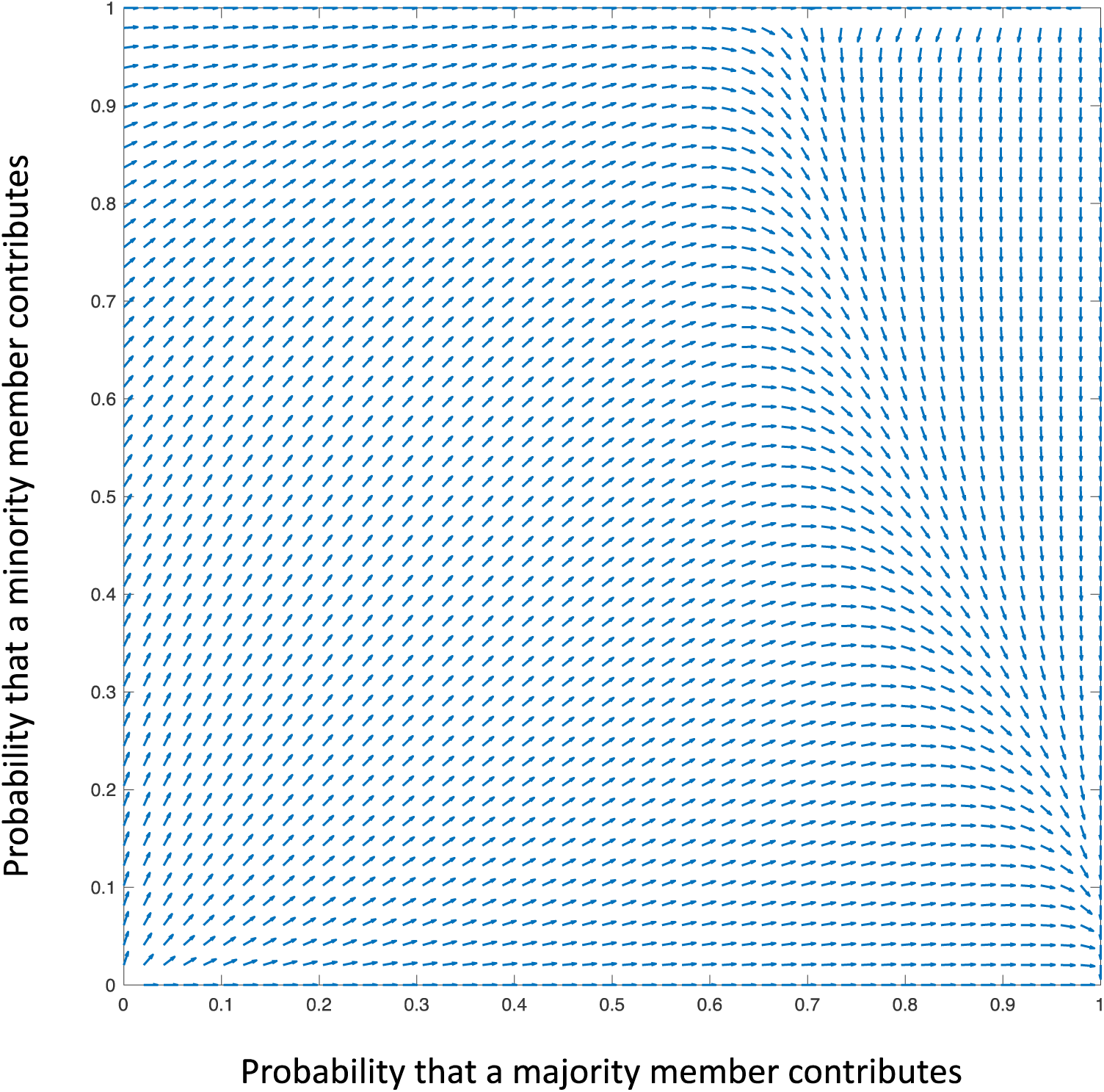
Evolutionary dynamics of the toy model with α = 1.1 (freeriders enjoy 10% higher fitness among the survived individuals, whether they are from the minority or the majority population). Vectors depict Δx, Δy (see Supplementary material) but normalized to be of equal length for clarity. Contributing is selected for (arrows point upwards and to the right) when it is only moderately present in either population, but once contributing becomes common, the minority is selected to decrease its contributions and the system evolves to x = 1, y = 0 where all contributions are produced by the majority.

We next proceed beyond the toy model, as real biological groups are rarely formed of two reproductively isolated types. (Mutualisms are an example, but they differ from our context because the two species involved typically do not contribute different quantities of the same resource — e.g. in rhizobia-plant mutualisms, plants do not contribute to nitrogen-fixing but instead offer other resources.) One context where conspecifics may consist of distinct categories, with the possibility of one being in a minority, is that of two sexes, with the potential for sex-specific selection to contribute to a task that could be performed by either sex in principle. In the context of avian cooperative breeding, there is support for the idea that adult sex ratio biases (which can arise through differential mortality and thus do not require primary sex ratio biases) can make the majority sex adopt the helper role (7). Here, difficulties in mate finding by the surplus sex play a considerable role in the relative profitability of philopatric helping as an alternative to independent breeding (see (8) for a related idea where sex ratios also evolve — the helper repayment hypothesis). In arthropods, selection to overproduce the more helpful sex can lead to the coevolution of helping and sex ratio (9, 10). To highlight differences between these earlier ideas and ours (while noting they are not mutually exclusive), we focus here on a setting that differs crucially from those considered in earlier work. In collective defence against predators by haplodiploid larval groups, mate availability is irrelevant (at this life-history stage), offspring do not interact with their mothers, and sex ratio biases arise easily because unfertilized eggs develop into males. Virgin females can thus produce sons, and mated females can adjust the sex ratio among their progeny by laying a certain proportion of unfertilized eggs.

Finally, we explore experimentally the effect of sex-ratio and kin-structure on the contribution to a collective act in larval groups of the haplodiploid pine sawfly *Diprion pini.* Similar to other gregariously living pine sawfly larvae, *D. pini* feed in dense groups during the larval stage (11). When threatened by natural enemies, larvae defend by regurgitating a resinous droplet of fluid from their mouth and perform a defensive display in concert (12). The sticky physical barrier produced by a group of defending pine sawfly larvae makes them unprofitable as prey for both avian and arthropod predators, as it is difficult for a predator to catch an individual without smearing its feathers or cuticle with the resinous fluid (12–14). Larval survival is consequently higher in groups than for solitary individuals (11, 15, 16).

Group defence in chemically defended prey can be considered to be a collective good, as (i) the survival of each individual within the group against predation increases with the proportion of defending individuals within its group (Lindstedt, Valkonen, Mappes *in prep*.) and (ii) contributing to group defence is costly for the prey (17). In pine sawfly larvae, individuals who contribute more to defence suffer decreased growth rates, lower immune responses and weaker chemical defence in future encounters (18). Sex-ratios of sawfly populations, including many pine-sawfly species, are often female-biased (19) and *D. pini* males in kin-groups are more likely to ‘cheat’ (fail to contribute to group defence) than their sisters (18). We performed two experiments testing for probability of showing defence behaviour in experimental attacks on an individual larva, to test if the proportion of larva that respond defensively per group (experiment 1) or the probability of an individual defending (experiment 2) depend on the group composition that the individual is from (sex ratio and kin vs. mixed).

## Results

### Model results

#### Sex ratio effect: Few males lead to females contributing relatively more

When larvae feed in kin groups, the primary sex ratio is a major determinant of the predicted sex-specific contribution to group defence (Fig. 2). If the proportion of males remains low, females contribute relatively more to group defence than males (and vice versa), but the magnitude of this effect is not identical across all low sex ratio situations. Collective action contributions are particularly female-biased (visible in Fig. 2 being reddest at the bottom right edge of the surface) when the rarity of males results from a low proportion of unmated females (low *p*_0_; unmated females produce sons only). The contribution bias remains milder when some females produce son-only broods while others specialize in producing females (higher *p*_0_, *r* < 0.5), i.e. conditions in which some broods lack females and males then need to defend without being able to expect contributions from sisters. As a whole, in line with the results from our toy model, the sex that is less common in the population is more prone to freeride. While the sex ratio effect is consistent across scenarios, sex-specific costs may modify the response in a logical direction (more male contributions if male reproductive success is not strongly hampered by defensive behaviour in the larval stage and similarly for females; compare columns in Fig. 2).

**Figure 2.**
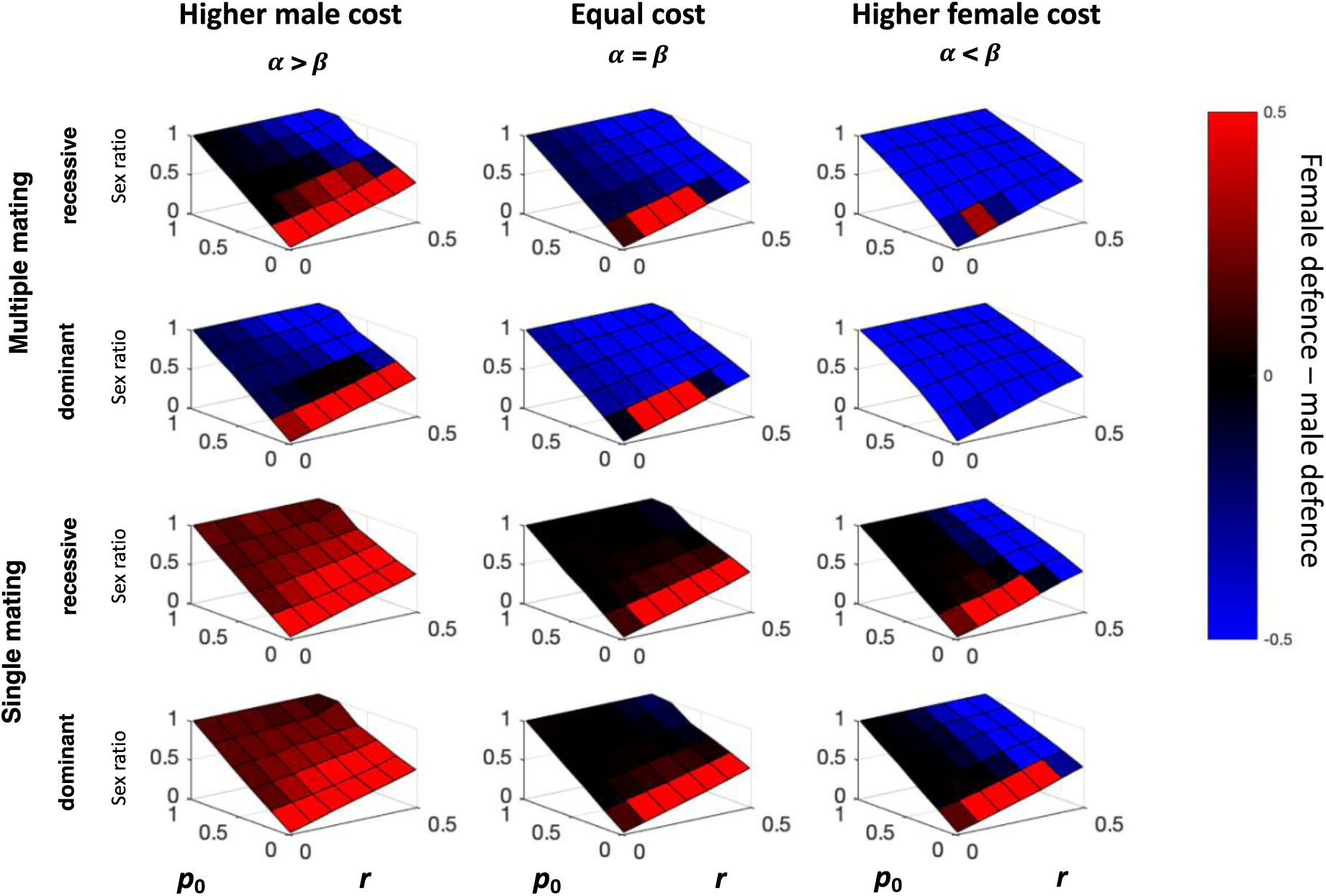
Mean male and female defensive behavior of 5 independent runs after 100 generations of simulation for varying *r* (x-axis) and *p*_0_ (y-axis) which together influence the sex ratio in the population (z-axis). The evolved relative contribution of females and males to the group defense is indicated by the coloration (female defense – male defense), with redder coloration indicating that females contribute more (per capita) and bluer coloration that males contribute more (per capita) to group defense. Female effort associates with female-biased sex ratios, male effort with male-biased sex ratios, but the borders between male- and female-biased effort are modulated by dominant vs. recessive gene expression and multiple mating (if permitted, females contribute less). Parameters: increased male cost, α= 1.5, β= 1.2; equal cost, α= 1.2, β= 1.2; increased female cost, α= 1.2, β= 1.5. Other assumptions: γ=3, kin-structured groups, and independent loci for male and female behavior.

#### Dominance and multiple mating increase female investment in group defence

Under haplodiploidy, males are bound to express their defence alleles, while in females, defence alleles may be impacted by dominance. This turns out to only have a strong effect in mixed groups when costs of defending do not differ between the sexes (Fig. S1, middle column), or alternatively when we assume unresolved sexual conflict by making sex-specific expression constrained such that there is only one shared locus for defense behavior for males and females (with dominance assumptions impacting female but not male behavior (Fig. S2-S3). Where dominance has an effect, it makes females contribute relatively more.

In kin groups, multiple mating has a much clearer effect than dominance. It weakens female relatedness with no effect on males (as they are fatherless), and the consequence is in the expected direction: the relative contribution by females decreases (making the top two rows in Fig. 2 much more ‘blue’ overall when multiple mating is assumed). As expected, this effect is not driven by an increase in absolute male contribution, but rather stems from a decreased contribution by females (absolute values shown in Fig. S4: females contribute less in scenarios with multiple mating).

#### The sex-specificity of the contributions disappears if contributions from everyone are essential

Our main results were derived with *γ* = 3, implying that group survival remains relatively intact across a low to moderate proportion of freeriders (2^nd^ column of Fig. S5). Higher values of γ imply that group survival can tolerate an even higher proportion of freeriders; lower values mean that even if one individual fails to contribute, this is ‘felt’ by the entire group (left column of Fig. S5: survival drops immediately if some freeride). Intuitively, one would expect that if everyone’s contributions are essential, all will contribute and the sex bias disappears; the results support this (Fig. S5, sex biases require a large enough *γ*; the sex bias first appears at low sex ratios, as in our main results, and with a high enough *γ* spread to all sex ratios).

### Experimental Results

We estimated the influence of larval group structure (kin vs. mixed groups), sex ratio as well as the absolute number of males and females on plasticity in defensive behavior by fitting a set of candidate models to the experimental data (response variable: proportion of individuals in a group that display defensive behavior when experimentally attacked) and estimating their support based on Akaike information criteria (AIC). The ‘attack’ was performed by gently squeezing a randomly chosen larva within a group with forceps. The best-performing model included sex ratio and group structure (kin vs. mixed groups), and predicts that (i) the number of individuals displaying defensive behavior decreases with the number of males, and (ii) individuals in kin-groups contribute more to group defense than those in mixed groups (Table 1a, Fig. 3a&b).

**Figure 3.**
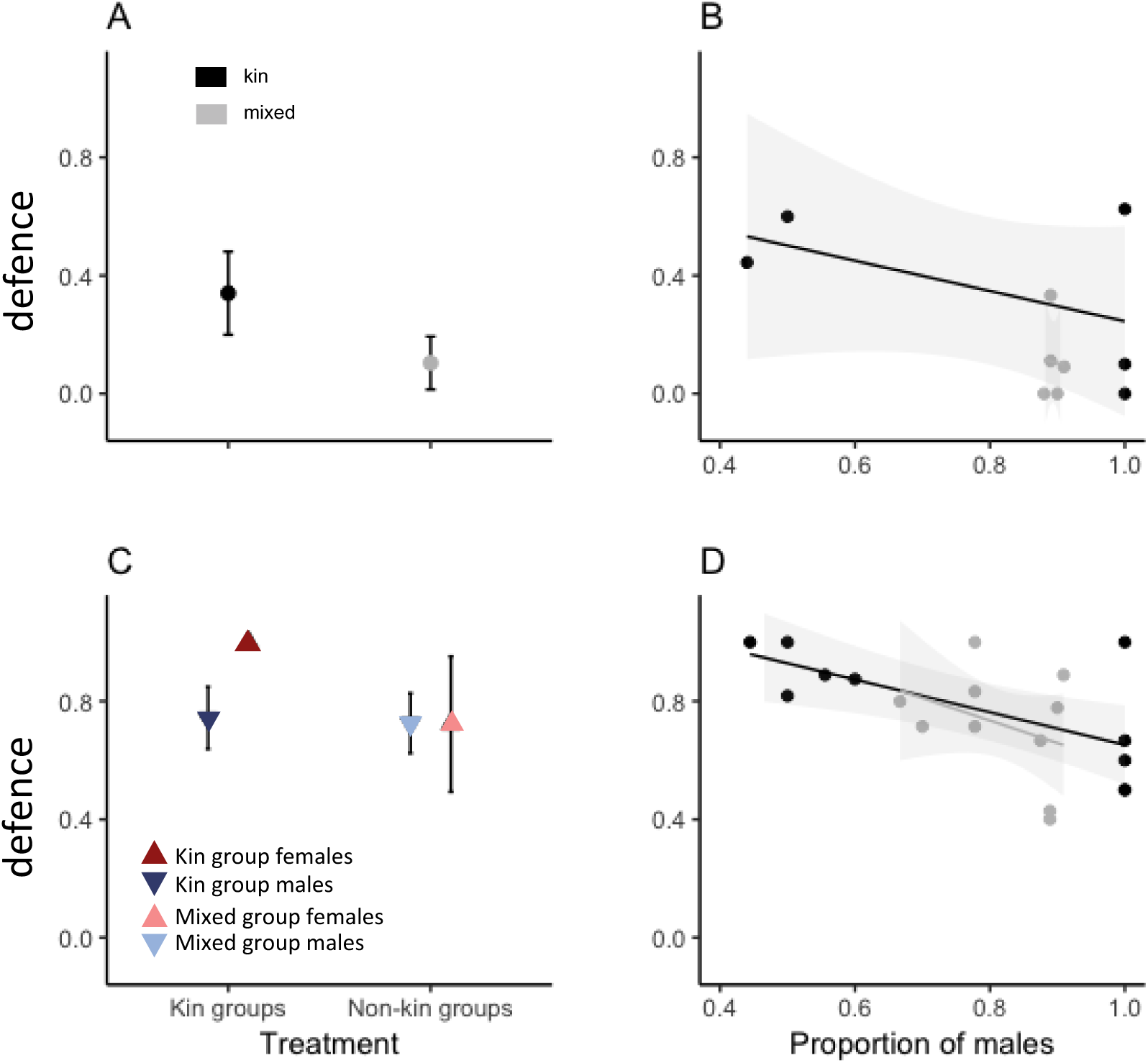
Experimentally determined proportions of defending individuals in entire groups (A,B) or when tested individually (C,D). The same data are shown both with respect to treatment (kin groups vs. mixed groups, A, C) and against the proportion of males in the groups that the larva came from (B, D). Means of the proportion of defending individuals are shown with confidence intervals.

**Table 1.** The tested models with the respective AIC, (Akaike Information Criterion) scores. Models are listed in order of increasing AIC values (AIC, score differences ΔAIC) for the group data (a) and individual data (b). Green shading shows models within 2 AIC points of the best model.

| Model predictors for proportion of defending individuals (kindata) | AIC | ΔAIC |
| --- | --- | --- |
| sex ratio and kin structure | 337.163 | 0.00 |
| number of females and kin structure | 337.229 | 0.07 |
| kin structure | 337.986 | 0.82 |
| number of males and kin structure | 339.005 | 1.84 |
| number of males and number of females and kin structure | 339.039 | 1.88 |
| number of females | 373.307 | 36.14 |
| null model | 381.917 | 44.75 |
| sex ratio | 383.878 | 46.71 |
| Model predictors for proportion of defending individuals (individual data) |  |  |
| sex ratio | 132.406 | 0 |
| own sex and sex ratio | 133.744 | 1.34 |
| sex ratio and kin structure | 134.158 | 1.75 |
| own sex and sex ratio and kin structure | 135.433 | 3.03 |
| own sex and sex ratio and kin structure and random number | 137.370 | 4.96 |
| own sex | 138.166 | 5.76 |
| own sex and kin structure | 138.408 | 6.00 |
| kin structure | 139.644 | 7.24 |
| null model | 140.001 | 7.60 |
| sex ratio | 132.406 | 0 |

The above results (Table 1a) are based on data collected at a stage where the sex of the larvae cannot be evaluated: larvae were only sexed after pupation, i.e. the experimenter was blind to group sex ratios at the time of the experiment. This experiment therefore does not comment on sex-specific investment directly, as it yields no direct measurements of the proportion of male and female larvae that responded when a sister or a brother was attacked. However, the group sex ratio was known afterwards, and both the best-performing and the second-best model in terms of AIC (the 2^nd^ has roughly equal support ΔAIC=0.07 as the first) include a sex ratio effect: the best model predicts that the proportion of defenders increases at female-biased sex ratios, the 2^nd^ model yields the same outcome by specifically including the number of females (instead of sex ratio), as well as group structure, as predictors of the proportion that defend. The third model with good support includes group structure with no other predictor (ΔAIC=0.82). Two other candidate models are within the range of two AIC points. Both include the number of males (which reduce contributions to defense) and group structure, and one additionally includes number of females (which increase contributions) as a predictor (see Table 1a). Overall, these data are consistent with the idea that female larvae, though unsexed at the time, were responsible for producing the majority of defense, and that larvae defended more if surrounded by kin (Table 1a, Fig. 3a&b).

We also performed an experiment (Table 1b) where larvae of the last instar were attacked individually, recording whether they deployed the defensive fluid. Since the larvae were kept individually after this experiment, the sex could be assigned retrospectively once the larvae pupated (20). All models with at least moderate support (within ΔAIC < 2) include the sex ratio of the group that the larva came from as a predictor (Table 1b, Fig. 3 c&d), with males defending relatively more in male-biased experimental groups and females in female-biased ones. The differences between the three well-supported models relate to whether own sex, group structure, or neither is included in addition to group sex ratio (Table 1). In models where they are, being female increases defence probability, similarly for being in a kin group.

## Discussion

We find mathematical and experimental support for our hypothesis that when groups consist of demographically distinct roles (different sexes in this case), that are not represented equally in the population, the majority has a tendency to evolve to contribute more to the collective good. The minority is more prone to freeride, as its failure to contribute to the collective good does not have as severe consequences for the entire group than if the individuals forming the majority did so.

Our toy model confirms this logic in the simplest possible setting, while our full model is based on a real-life example involving haplodiploidy and a trait that is expressed long before reproductive activities and the associated sex-specific behaviours begin. In the life cycle of the full model, multiple other asymmetries than that of a ‘minority sex’ – ‘majority sex’ may also play a role. Specifically, female relatedness may vary as a result of multiple mating while that of males does not. This can also modulate how much females contribute; less in case of multiply mated mothers producing kin-grouped larvae than if the mother was singly mated. Additionally, female (but not male) behaviour is impacted by a diploid genome, with the potential for dominance to act on gene expression. However, this aspect of haplodiploidy is not the main driver of our results: dominance only had a marked effect in model variants that assumed females and males cannot evolve their contributions independently (i.e. unresolved sexual conflict). In this case recessivity, rather obviously, reduces the probability that a female expresses the trait, for any frequency of the trait in the population.

Our model also predicts group structure to play a role. The result that there is more defence as a whole in kin-groups than in mixed groups is unsurprising. It is more intriguing that the way sex ratio biases come about can have an impact on the sex-bias in behaviour. If male-only broods increase in frequency (because some mothers reproduce as virgins), males contribute more to defence than if a similar sex ratio is produced through primary sex ratio adjustments among mated mothers. In the former case, male larvae are more likely to exist without any sisters, and lack of defence would be very detrimental to such groups.

Many of these predictions are borne out by our experimental data from the haplodiploid *D. pini* larvae. Individuals contributed more for the collective larval group defence in kin-groups compared to mixed groups (in which individuals originated from different colonies), and although an AIC analysis could not unambiguously differentiate between all variant models in which own sex, group sex ratio and group structure (kin or mixed) were included or excluded, the overall message remains clear: the more females in a group, the higher the proportion of individuals contributing to defence.

Our mathematical model makes a number of key assumptions. Ours is an ‘open model’ (sensu (21)) where some parameters, notably the probability that a female is unmated and the sex ratio produced by the mated ones, are evaluated at every possible combination, rather than restricting the view to those combinations that are more likely to be found in nature. In reality, if many sons will be produced by virgin mothers, there will be selection on mated mothers to specialize in the production of females, more so than they would in the absence of virgin reproducers (22). Effectively, this means that the most likely areas of our surface plots found in nature are those that do not feature unrealistically high proportions of males. We still prefer to show the outcomes (male-biased contributions to the collective good) for those regions too, as this helps to understand what conditions would be required for those to happen, and strengthens the general message: the majority-minority asymmetry has the main effect that we report, regardless of whether the majority consists of diploid or haploid individuals.

Second, we assumed a rather stark contrast between completely resolved vs. completely unresolved sexual conflict. Larval defence behaviour in our models was either determined by completely different loci controlling the same behavior in male and female larvae, allowing a resolution of the conflict, or a single locus being expressed in both males and females (though in a diploid state in the latter, in which case we investigated complete dominance and recessivity as the two extremes). Here, our aim was to chart the extremes between which real cases, with possibly polygenic expression of traits, are likely to lie. Completely independent loci allow each sex to approach its own optimum in a manner that is dependent on what siblings of a different sex *do*, unencumbered by genetic correlations between brother and sister loci. The genetic architecture underlying defensive behaviour in *D. pini* is currently unknown, but in general *D. pini*’s likelihood to cheat in defence in larval stage (not produce defensive secretion under attack) is a partly heritable trait both in females and males (18).

In our model, the sex-ratio effects were stronger in kin-groups than in mixed groups. In general, genetic relatedness promotes cooperation and increases contributions to collective action. For examples, both theoretical and empirical research on the evolution of cooperative breeding in haplodiploids suggest that cooperation is more likely to evolve under monogamy than when broods are the result of polyandrous matings (23–26); in theoretical work that specifically asks whether males or females should evolve into helpers, monogamous situations under haplodiploidy can favour females as helpers (9, 27). Our results add a collective action angle to these results: ours is not a model of cooperative breeding, as in our setting no individual foregoes own reproduction to help others, instead the collective good is group survival up to maturity. It also highlights a general principle about minority and majority roles; while coexistence of cooperators and defectors is possible without such subdivision (1), our results show that limits to the frequency of freeriders that arise from population subdivision can help stabilize cooperation in the presence of cheaters.

Our new experimental results corroborate the findings of earlier research (18), suggesting that *D. pini* females were more likely to contribute to group defence in kin-groups than males. In our current study, we found evidence that the sex-ratio of the group modulated larval defence behaviour. Although our study design made it difficult to disentangle whether female-biased groups cooperated more simply because they had more females (that each contribute more) or whether males too increase their effort in the presence of many sisters, our results raise the possibility that *D. pini* larvae have some means to recognize their kin as well as the sex of the other individuals in the group, adjusting their cooperative behaviour accordingly. Potential mechanisms include CHP-compounds, already known to be important in nest-mate recognition both in immature and mature life-stages in eusocial ant species (28, 29). In this context, it is interesting to note that individuals were divided into the experimental groups as smaller instars, and differences in their defensive behaviour were consistent until the last larval instar. Cues to recognize kin therefore appear to be either innate or very strongly impacted by very early natal environment (before we mixed the larvae).

Our experimental data was in one sense non-ideal for testing our results: natural sex-ratios in sawflies that tend to be female biased (19), but some of our mothers, despite mating with a male, only produced male offspring, i.e. we may have been testing our predictions under a larger *p*_0_ value than is the naturally expected norm. Since we could not determine the sex of the larvae until pupation, using all-male larval groups resulted in male-biased sex-ratios, which impacted the sex ratios that were created for the mixed-group treatment. Despite the generally low proportion of females in such groups, the outcomes were robustly in line with our model predictions, which fortunately are directional (more females leads to more total contributions to the collective good, whether the baseline sex ratio is low or high).

Our model considered contribution to collective antipredator defence in a specific setting, group living haplodiploid larvae, but may open up interesting avenues for future research to study effects of sex-linked differences in collective traits in general. Ours is not the only species where females appear to contribute more to chemical defence: in burying beetles, females can deploy higher quantities of the defensive secretion (30), while in other systems females concentrate more toxins in their bodies (31), or transfer toxins for their offspring (32). Future studies could usefully test the extent to which these reflect life history investment differences that are selected for at the individual level, or are potentially shaped by selection to maintain a collective good via educating the relevant predators. There are also interspecific settings where rarity of one type is not related to abundances of the two sexes, but of two (or more) species. As a concrete example, public goods games exist between species in aposematic chemically defended insects that share the costs of predator education (predators learn to associate warning coloration with the chemical defence of the prey and avoid attacking them and any similar species in future). If defence is costly for an individual, some may ‘cheat’ by relying on the already existing protection rather than contributing to predator education ((17); for a multi-species setting where larvae are also taking advantage of the temporal order offered by seasonality, see (33)). More generally still, the categories of individuals involved may also be sexual and asexual forms of the same species, (34), or cell types in cancer, (35), highlighting the general need to understand heterogeneity of individual contributions in the light of the demography producing these individuals.

## Methods (See SI for details)

### Model overview

We created individual-based simulations of a population of haplodiploids with non-overlapping generations. Haplodiploidy often leads to biased sex ratios e.g. (9, 26, 36, 37), and we model the larval sex ratios as a result of two processes: some females (proportion *p*_0_) remain unmated and produce only male offspring, while mated females can produce a mix of male and female offspring; we model their sex ratio as *r*. We vary *p*_0_ from 0 to 1 and *r* from 0 to 0.5, but do not let them evolve, as our aim is to examine the consequences of biased larval sex ratios on larval contributions to a common good, rather than examine the evolutionary reasons for sex ratio biases per se.

We examine 2 x 2 x 2 x 2 different scenarios (Fig. S8): (i) larvae can either stay in their original kin groups or mix into an equivalent-sized but unrelated group, (ii) the mother of the larval group has mated multiply or singly (in case she has mated at all; in either case we assume that a proportion *p*_0_ of females do not mate at all), (iii) the contributions may be dominant or recessive when the individual is diploid (female), thus phenotypically, each individual is a contributor or free-rider, and (iv) the cooperative behaviours is determined by independent loci for females and males or by just one shared locus. In the independent locus scenario, to allow male and female behaviour to evolve separately, we model the evolutionary dynamics of 2 loci (haploid in males, diploid in females) that determine whether they invest in group defence or not; note that this is not fundamentally different from an alternative approach where the same loci determine a baseline while another locus changes the expression based on an individual’s sex (either way, we allow sex to impact behaviour if a sex difference is selected for); Thus females have two copies of the male defence allele but do not express either of them, and female expression of their own defence alleles can be either dominant or recessive. Males also carry one copy of the female defence allele but do not express it. In the scenario with the same locus for males and females, females still carry 2 defence alleles and expression can be dominant or recessive, whereas males only carry one copy of the defensive allele.

The model traces stochastic evolutionary dynamics when group survival is a declining function of the proportion of free-riders with a parameter *γ* describing how well survival is kept intact even if freeriding is of appreciable frequency (see Fig. S6, full details are given in Supplementary material), and runs until the proportion of free-riders and contributors stabilizes. In all figures, we show the mean of 5 independent runs after 100 generations (Fig. S7). Adult individuals that were phenotypic free-riders during their larval time have a sex-specific fitness advantage (α for males, β for females). A new generation is formed by sampling mothers and fathers among the survivors, taking into account α and β to make free-riders more prevalent as parents than their representation in the parental generation.

### Experimental Study

#### Study species

All *D. pini* families used in this experiment were from the fifth generation descending from a sample of 100 females and 100 males originating from an outbred laboratory population in the University of Berlin, Germany and reared on Scotch pine (*Pinus sylvestris*). In order to prevent inbreeding, the source population has been regularly supplemented with wild individuals collected from Germany.

After mating, *D. pini* females were allowed to lay their eggs on randomly chosen Scots pine branches. After the larvae hatched, they were fed with pine branches *ad libitum*, and fresh branches were provided twice a week. Throughout the experiment, larvae were reared under constant temperature (20 ± 2 °C) in a laboratory room which included both natural and electric light more or less constantly throughout the day. For the first 10 days after hatching, larvae were reared in family groups. After that, larvae were randomly divided for the treatments. In the experiment, we only used offspring of the females that were offered male to mate with and therefore were likely to produce sexually produced families. However, 4 of the females reproduced asexually and laid only haploid male eggs. Therefore, in non-kin groups, sex-ratios were male-biased (see below) and four of the kin-groups were all male-groups.

#### Experiment: Defensive behaviour in kin- and non-kin groups

To investigate if socially defending pine sawfly larvae are more likely to direct costly defensive acts to increase survival of their kin, we tested how individuals contributed to the cooperative defensive behaviour when they were reared in kin-groups (individuals originate from the same sexually produced clutch) or in non-kin groups. To control the heritable variation in the chemical defence (18), same families were split across the kin- and non-kin treatments. We had two blocks of 5 families that had all hatched on the same day. This was to ensure that larvae had similar size and developmental stage/instar (in case it somehow affects their cooperative behaviour). In the first block, those same 5 families were used to form kin-groups (all the individuals originated from the same family) and non-kin groups of 10 individuals (2 individuals per group originated from the same family). Thus, altogether we had 10 families and 10 kin-groups and 10 non-kin groups. Larvae were divided into treatments randomly. Since we cannot tell apart the sex of the individual in the early instars, we were not able to control the sex-ratios within larval groups and among treatments.

Defensive behaviour of the larvae was measured 3 times during the experiment: the first time 2 days after the experiment started, the second time after 7 days experiment started and last time when larvae were 21 days old before the final dispersing instar. All the individuals within one block (5 kin-groups and 5 non-kin groups) were measured on the same day. In the first two defensive behaviour assays (2 and 7 days), we measured the number of defending individuals per group when one randomly chosen individuals from the group was ‘attacked’ artificially by poking it 1-2 times on the dorsal side with the tip of the forceps. When larvae were 21-23 days old, we attacked the larvae individually by gently gripping them with the forceps and we measured the volume of the defensive fluid they produced as well as the size of the individual (length). In these measurements, the defence droplet was sucked into a 5μl capillary and its quantity was measured with an electronic ruler. We measured the body length of larvae with a ruler at the same time to control for the effect of body size on the amount of defence fluid produced. After defensive measurements, individuals were moved on Petri dishes to be reared individually to get individual information. Thus, the last round of defensive assays is the only one where we know the identity of individual such as sex. In addition, for part of the individuals (N =32), we lost the information of the defensive behaviour in the age of 21 days as they had already changed into final dispersing instar or pupated. These were all males as males pupate earlier. During the pupal stage, sexes were distinguished by pupal weight (20): individuals weighing over 100 mg were classified as females (mean pupal mass for eclosed females 127.71 ± 12.595) and below 100 mg as males (mean pupal mass for enclosed males 57.59 ± 9.368).

#### AIC Analysis

We estimated the influence of group structure (kin vs. mixed groups), sex ratio as well as the number of males/females on defensive behavior by fitting a set of candidate models to the experimental data and estimating their support based on Akaike information criteria (AIC) for both group and individual data. Each candidate model specifies the predicted probability to contribute to the group defense based on a multinomial logistic regression. Different models vary in the amount of parameter and the predictor variables. The tested models range from a null model, where we assume the same predicted probabilities applies to everyone (Table 1a, Model A), to models where the group structure (Kin groups vs. mixed groups) as well as the sex ratio, either as the relative proportion of males or the absolute number of males and females were included as predictors (Table 1a, Model H). The individual data allowed us to include the sex of the focal larvae (own sex) and thus the most complicated model included sex ratio, group structure and the own sex (Table 1b, Model I). AIC scores were then calculated based on the parameter values that maximized the likelihood of observing the pattern of binary contributing to group defense. Thus, the log Likelihood of each model was then calculated as the sum of the predicted probability to contribute of the larvae that actually do so plus the sum of the 1-predicted probabilities of the ones that don’t contribute: 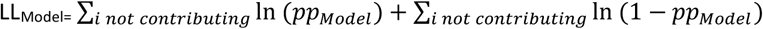. The AIC scores were then calculated based on the best total log-likelihood within each model, while penalizing models for including many parameters.

## Acknowledgements

This study was funded by the Academy of Finland via the project no 257581 (C.L.). N.G. and H.K. were funded by the SNSF via the project P2ZHP3_181500, 31003A_163374 and Finnish Centre of Excellence in Biological Interactions Research) grant no. SA-252411.

## Supplementary information

### Toy model of a collective good with two categories of players

Consider two large populations that are reproductively isolated but interact in the sense that, prior to reproduction, both populations contribute members to groups that participate in a common goods game. Population M (the majority) always contributes 3 members, population m (the minority) always contributes 2 members, such that the groups consist of 5 members in total. M-individuals contribute towards the common good with probability *x* or freeride (fail to contribute) with probability 1–*x*; for m-individuals the corresponding probabilities are *y* and 1–*y*.

We assume that if the group has more contributing individuals than freeriders, it will survive. The overall probability that a group survives is given by

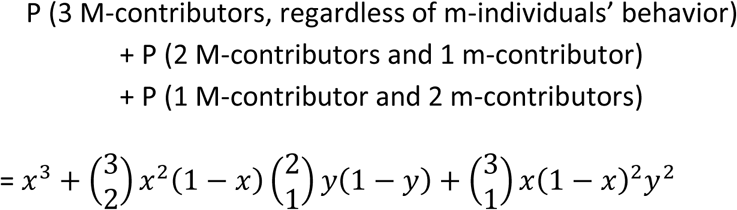

To evaluate evolutionary changes, we need to derive the fitness of contributing and non-contributing M- and m-individuals in a population where current contribution probabilities are *x* and *y*. We assume that if the group survives, the freeriders (who all survive) enjoy fitness α > 1, while surviving contributors have fitness 1. This reflects the costs of contributing in a common good game.

#### Fitness of a non-contributing M-individual (*W*_M0_)

Fitness is α for survived non-contributors, and survival depends probabilistically on the actions of other group members: survival occurs with probability

P(the other two M-members of the group contribute and at least 1 of the m-members does so] + P(one of the other two M-members contribute and 2 m-members do).

Thus

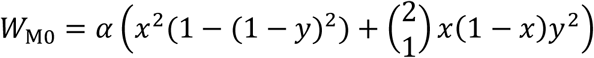

#### Fitness of a contributing M-individual (*W*_M1_)

Fitness is 1 if for survived contributors, and survival occurs with probability

P(both of the other M-members contribute, regardless of what the m-members do) + P(one of the other two M-members contributes and at least 1 m-member does so) + P(neither of the other two M-members contribute but 2 m-members do)

Thus

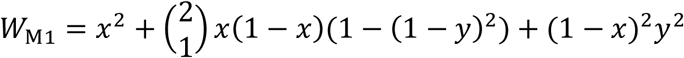

#### Fitness of a non-contributing M-individual (*W*_m0_)

Fitness is α for survived non-contributors, and survival occurs with probability

P(all three W-individuals contribute)

+ P (two of the W-individuals contribute, and the other w-individual does so too)

Thus

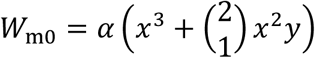

#### Fitness of a contributing M-individual (*W*_m1_)

Fitness is 1 conditional on survival, and survival occurs with probability

P(all 3 W-individuals contribute)

+ P(2 of the W-individuals contribute)

+ P(1 W-individal contributes and so does the other w-individual)

Thus

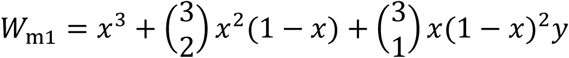

#### Allelic frequency change

Assuming asexual reproduction and genetically determined behaviour, we can calculate the changes over one generation as follows

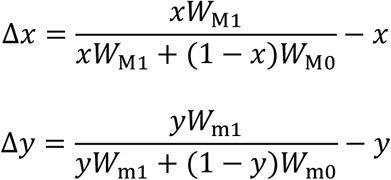

A strict demographic interpretation requires that each individual in each population participates in exactly 1 group formation event before reproducing; this is not incompatible with the assumption that populations are large, if the M-population as a whole is 50% larger than the m-population.

### Full haplodiploid model details

#### Initialising population

We initialised the population by creating *N* diploid individuals with either two loci for defence, one expressed in males and one in females, or just one shared locus (expressed in both sexes). We randomly assigned either 1 or 0 for each of the four or two potential alleles (2 for female and 2 for male larval defence if the latter are modelled separately). Thereafter, we convert (initially) half of the population into haploid males by retaining only the first of the two copies of the locus. To give maximal flexibility for the evolutionary process to find female and male optima without genetic constraints in the two-locus case, these loci are unlinked.

We begin the simulation with the adult generation (see below), however since the phenotype of each individual can be deduced from their genotypes, we assume these adults already have a larval past during which they were defenders or freeriders.

#### Life history events

##### Mating & reproduction

Adult individuals form one mating pool, and we first compute male and female competitiveness. When competing to be parents of the next generation, individuals that have been freeriders as larvae get a competitive advantage of *α* (for males) or *β* (for females). Since the competitive advantage is based on the phenotype, in the recessive scenario females only defend (and pay the cost of not obtaining the competitiveness benefit *β*) if they are homozygous.

Female competitiveness is expressed as the expected number of eggs they produce; females that were phenotypic freeriders as juveniles are produce a Poisson-distributed number of eggs with mean 10β, while phenotypic defender females’ number of eggs is 10, also Poisson-distributed. Additionally, and independently of competitiveness status, females differ in whether they have mated or not. Each female has a probability *p*_0_ of staying unmated; all the offspring of these females are male. Mated females use a sex ratio of *r*, interpreted as the probability that each of her eggs develops without being fertilized, i.e. becomes a male. Males inherit one of the two alleles of their mother (if two loci are inherited, inheritance is without linkage). Sexually produced offspring inherit one allele from their mother (similarly to the asexually produced males) and the second allele from a father.

All males compete to be chosen as fathers of mated females’ female offspring. For each fertilized egg, a male’s relative competitiveness is 1 if the male was a defender, and α if he was a freerider. In case of multiple mating, these competitiveness values determine the probability that a specific male is the sire for each fertilized egg independently of other eggs. In single-mating scenarios, the sire is chosen on a per-mother basis, with the same sire then used as a father for all female offspring for that mother.

Additional to their genotypes, offspring are assigned a group ID (identity of their mother), which is used to determine group survival, and optionally to shuffle individuals between groups, depending on the scenario under investigation.

#### From genotype to phenotype

We assume that all offspring experience attacks by predators, thus the defense genotype is expressed phenotypically as ‘freerider’ or ‘defender’ during the larval phase. Male offspring defend according to their haploid allele, females defend depending on assumptions we make on dominance. If defending is recessive, 00 and 01 females are freeriders, and 11 females defend. If defending is dominant, then 00 females are freeriders, while 01 heterozygotes as well as 11 homozygotes are defenders.

#### Group composition and survival

Groups are formed by either sticking with the brood structure that was obtained during reproduction (one group per mother), or by retaining the group-specific number of larvae while reshuffling all group identities of offspring (which destroys kin structure). Groups that are thus formed differ in their total number of individuals *n*, as well as the number of freeriders and defenders, which we denote *n*_f_ and *n*_d_, respectively.

Survival of larvae is determined in two consecutive steps. In the first step, entire groups survive or fail to do so, with survival

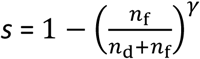

denoting the group-specific probability that offspring of a particular group have a chance to mature. After removing the individuals that fail in step 1, offspring may still succumb to other causes of mortality (step 2 of mortality), which we assume density-dependent with all offspring from all surviving groups pooled. If the number of potentially maturing offspring exceeds the carrying capacity *N*, we choose a total of *N* offspring to form the mating pool of adults in the next generation. In case fewer than *N* offspring survive step 1, then all of them are assumed to survive step 2.

#### Simulations

Simulations were performed for *t*_max_ = 100 years which allowed the mean male and female behaviour to stabilise (Fig. S6). We examine the results of these simulations assuming i) or recessive inheritance of defensive behaviour, ii) multiple mating or non-multiple mating iii) offspring remain in kin groups or form unrelated groups and iv) the cooperative behaviours is determined by independent loci for females and males or by just one shared locus (see Fig. S7). We contrast the equal cost scenario, where α=β, with dynamics where one sex benefits more from freeriding than the other. For all parameter values, we record phenotypic defence behaviour and the offspring sex ratios that are the result of varying *p*_0_ and *r* in each combination of the above scenarios, allowing us to test how haplodiploidy, sexual conflict and sex-ratio dynamics contribute to solving the collective goods problem in cooperative group defence.

## Supplementary Figures

**Figure S1.**
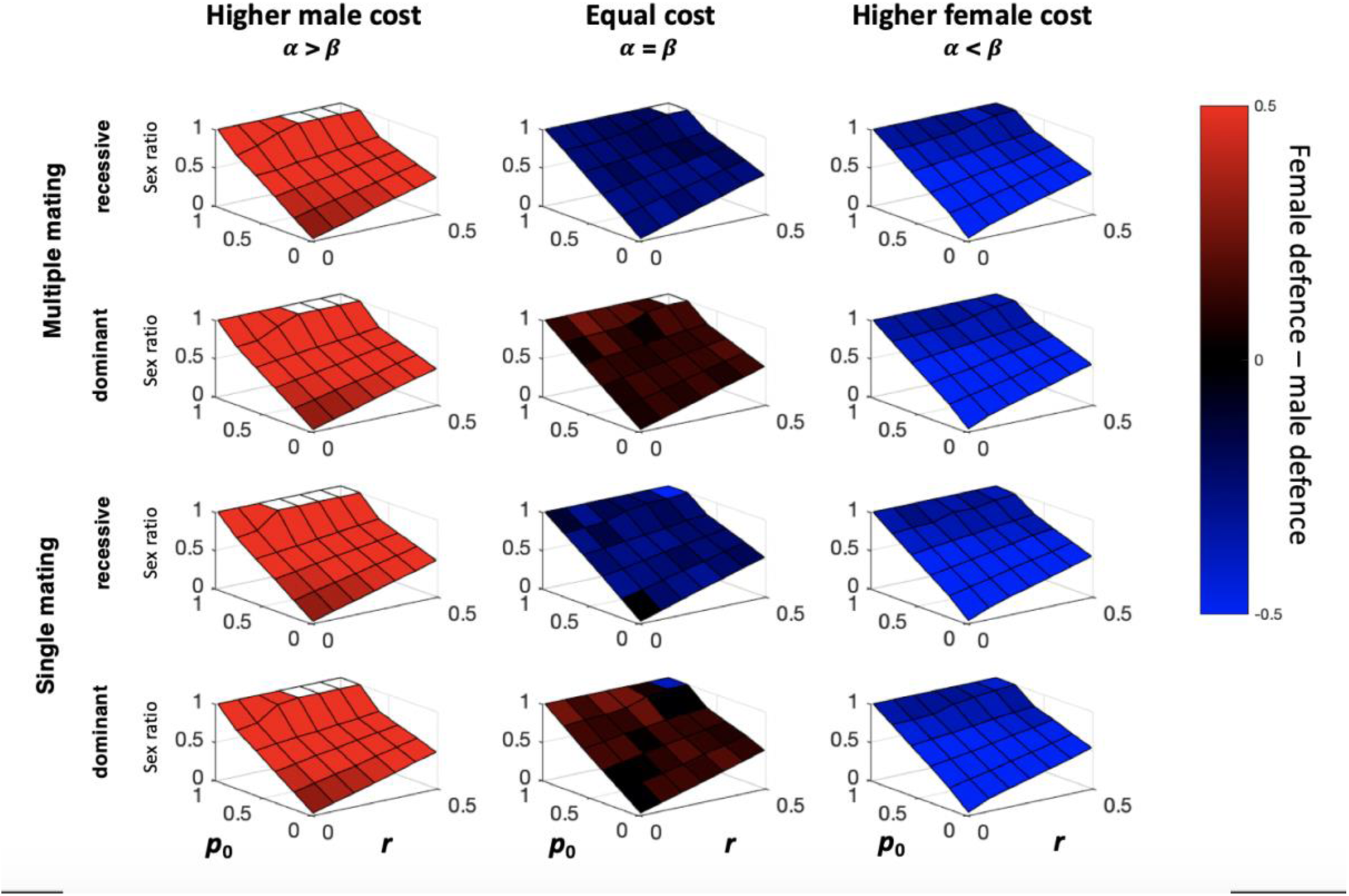
Same parameter settings as in Figure 2, but with a mixed group structure (no kin association). White squares indicate that populations went extinct and thus information on defensive behaviour is missing.

**Figure S2.**
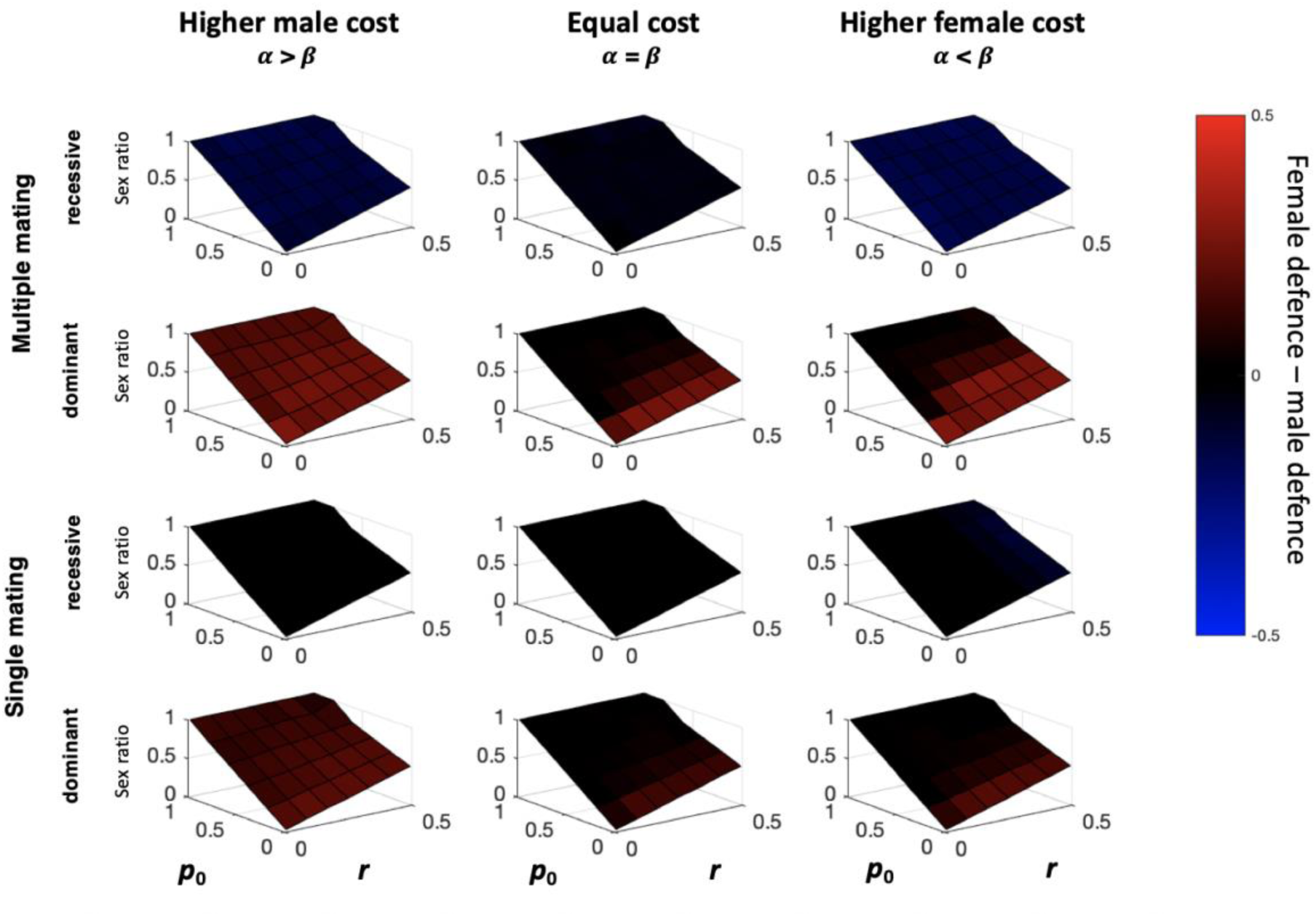
Same parameter settings as in Figure 2, but with one shared locus that determines both male and female behaviour.

**Figure S3.**
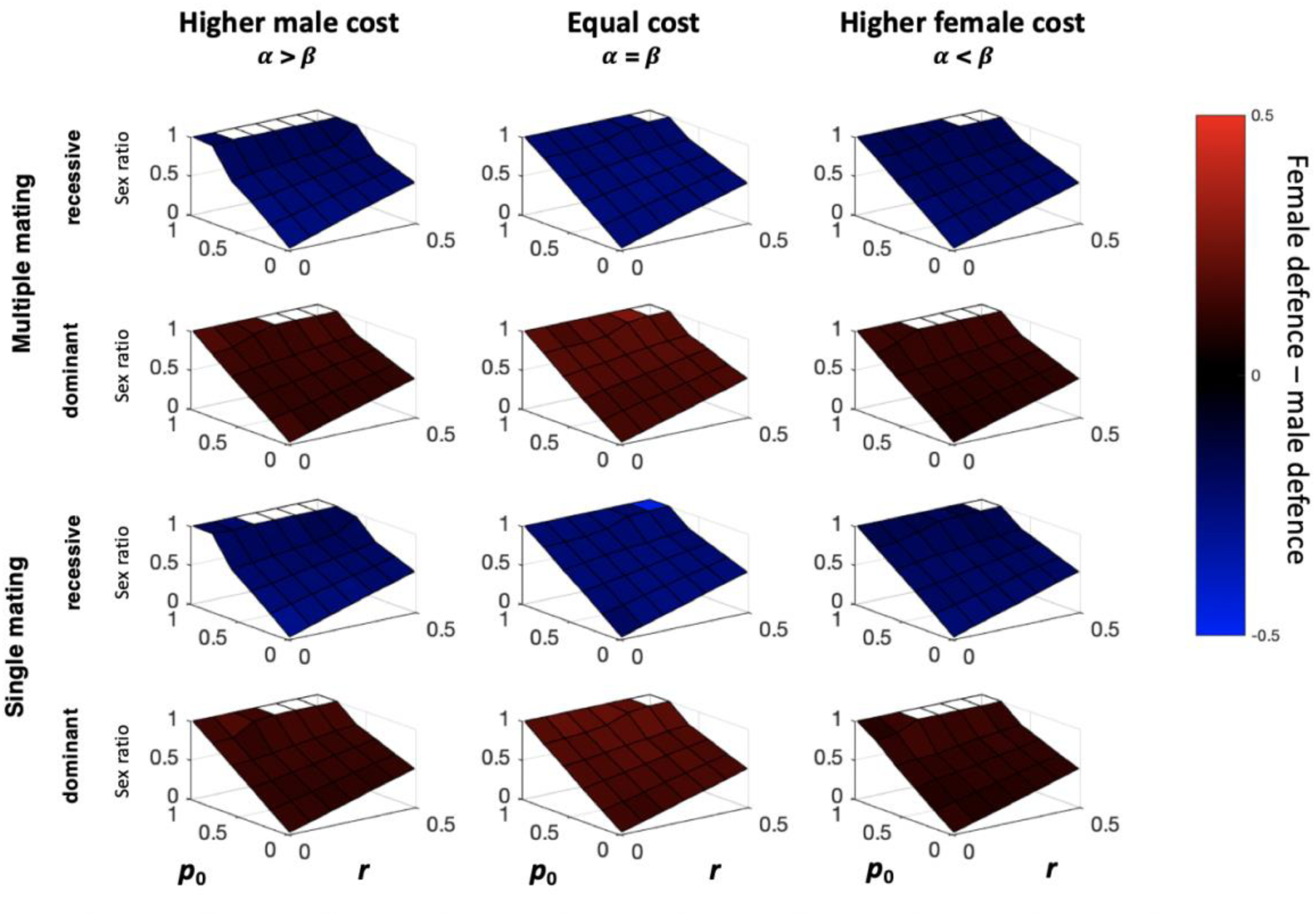
Same as Figure 2 but with one shared locus for male and female behaviour mixed groups (no kin association between larvae).

**Figure S4.**
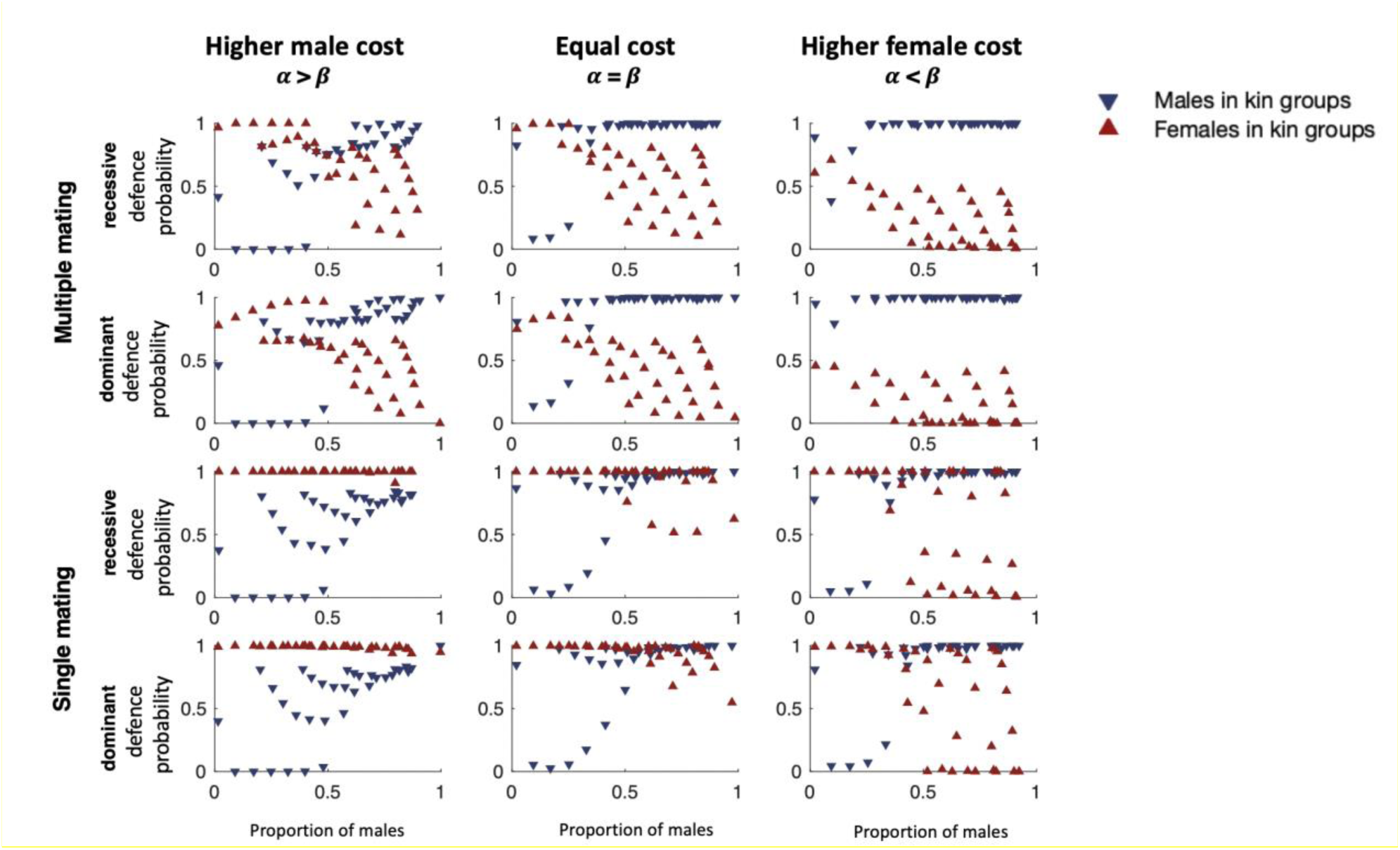
Mean male and female defensive behaviour in kin groups of 5 independent runs after 100 generations of simulation for varying *r* = {0.010, 0.108, 0.206, 0.304, 0.402, 0.5}, and *p0* ={0.0100, 0.1750, 0.340, 0.505, 0.670, 0.835, 1 which together influence the sex ratio in the population (placement along the x-axis). All subplots assume γ=3, and independent loci for female and male contributions, i.e. depicting absolute contributions for all cases across Fig. 2. Male contributions have the potential to exceed female contributions at high sex ratios and/or if defending is more costly for females.

**Figure S5.**
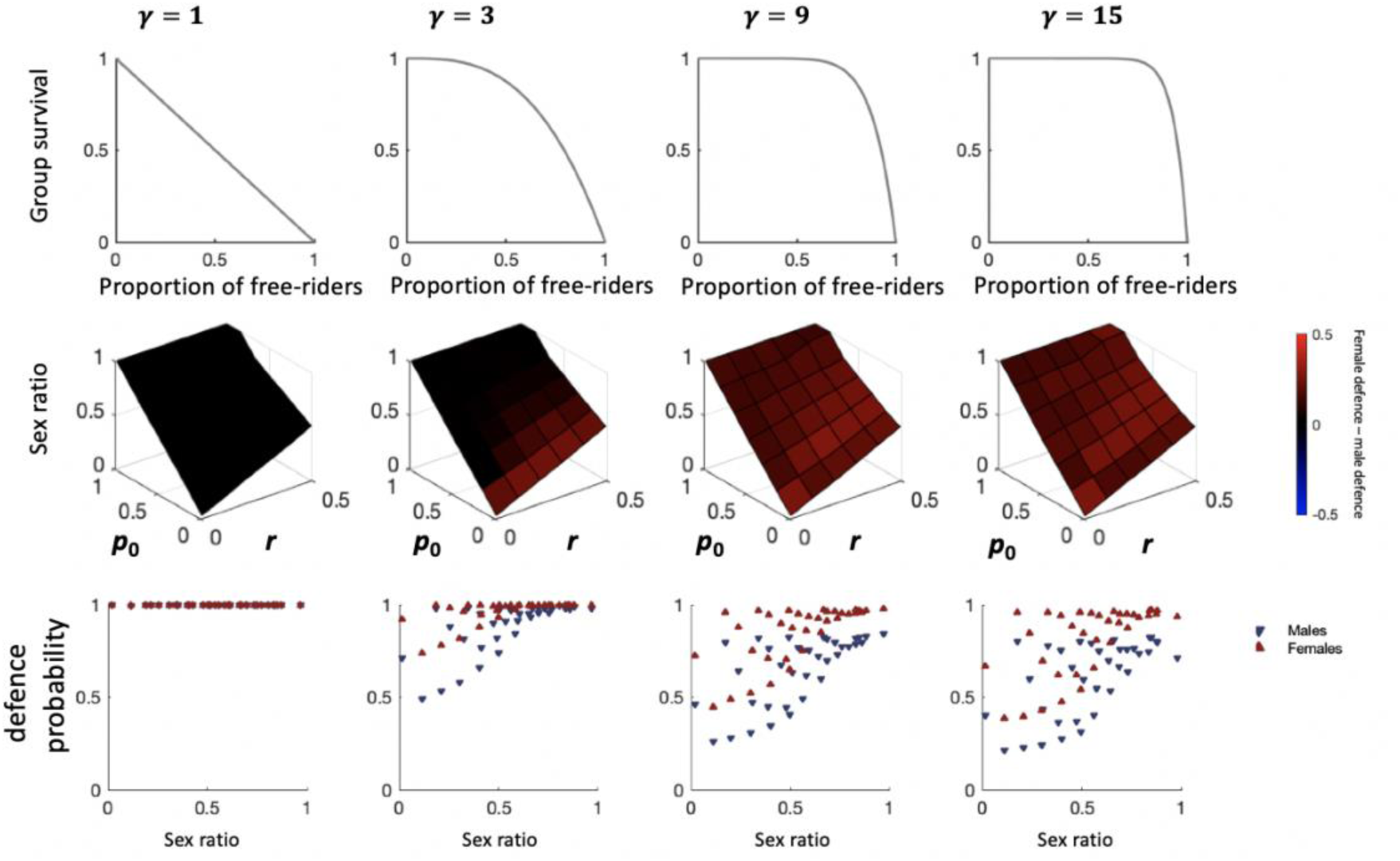
Effect of γ, a determinant of the relationship between the proportion of free-riders in the population and the group survival (top row), on the relative contributions of males and females (middle row, using the same colour scheme as in Fig. 2), and the absolute contributions plotted against the sex ratio that results from each combination of *p*_0_ and *r* (bottom row).All parameters and modelling choices, except the value of γ, are identical to values in Fig. 2.

**Figure S6.**
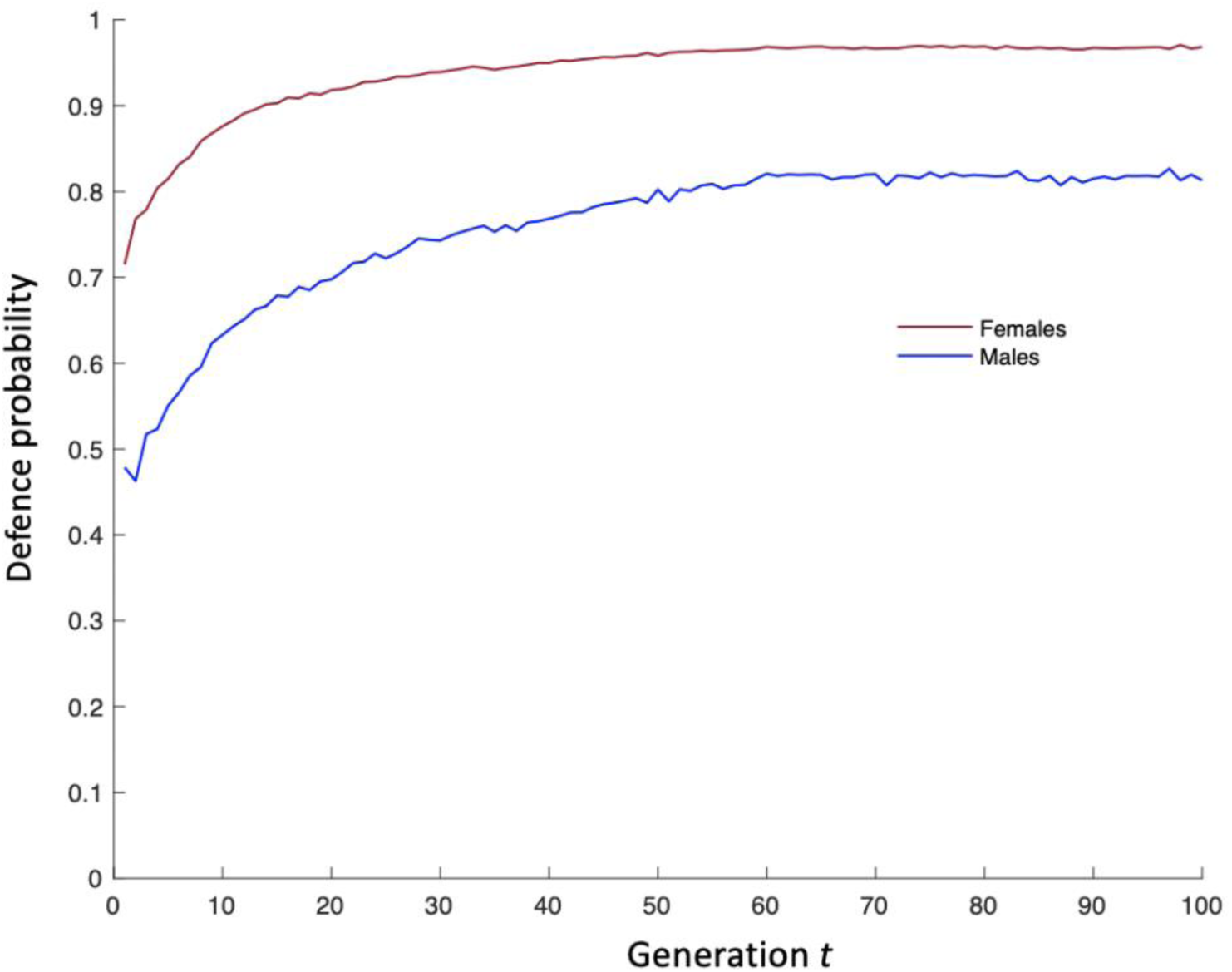
Mean probability of defence behaviour of male and female of 5 independent model runs over 100 generations. Dominance, kin groups, multiple mating, α= 1.2, β=1.2, γ=3, *r*=0.2, *p*_0_=0.2.

**Figure S7.**
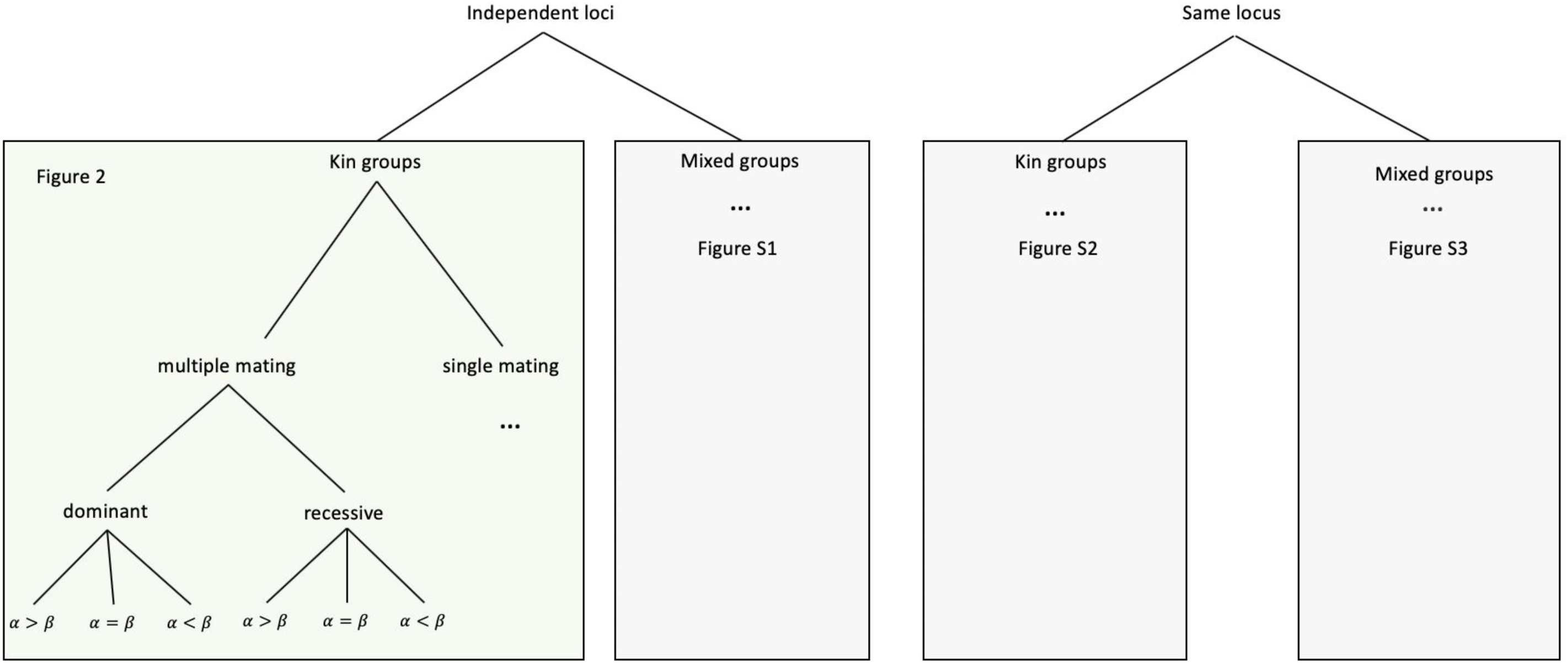
Overview of model scenarios and respective figures

